# Explainable Feature Extraction Using a Neural Network with non-Synaptic Memory for Hand-Written Digit Classification

**DOI:** 10.1101/2021.04.06.438597

**Authors:** Faramarz Faghihi, Hany Alashwal, Ahmed A. Moustafa

## Abstract

The human brain recognizes hand-written digits by extracting the features from a few training samples that compose the digit image including horizontal, vertical, and orthogonal lines as well as full or semi-circles. In this study, we present a novel brain-inspired method to extract such features from handwritten digits images in the MNIST database (Modified National Institute of Standards and Technology database). In this study, we developed an explainable feature extraction method for hand written digit classification in which the extracted information are stored inside the neurons as non-synaptic memory manner. For this purpose, a neural network with 10 single neurons was trained to extract features of training images (each neuron represents one digit class). Following that, the trained single neurons are used for the retrieval of information from test images in order to assign them to digit categories. The accuracy of the classification method of test set images is calculated for different number of training samples per digit. The method demonstrates 75 % accuracy using 0.016 % of the training data and maximally shows accuracy 86 % using one epoch of whole training data. The method as an understandable feature extraction method allows users to see how it works and why it does not perform well on some digit classes. To our knowledge, this is the first model that stores information inside single neurons (i.e., non-synaptic memory) instead of storing the information in synapses of connected layers. Due to enabling single neurons to compute individually, it is expected that such class of neural networks show higher performance compared to traditional neural networks used in complicated classification problems.

## Introduction

The accelerating progress in experimental and theoretical neurosciences has enabled Artificial Intelligence (AI) researchers to develop ‘Brain-Inspired neural networks and algorithms’ to overcome complicated AI problems [1, 2]. Although we are at the beginning of exploring the mysteries of neural computations, we know that neurons and small neural networks in different regions of the human brain are essential computational units. Experimental and theoretical studies have shown nonlinear calculations in synapses and the soma of neurons to integrate thousands of inputs from other neurons to generate spikes.

The human brain has been a source of inspiration in AI. Interestingly, on the other hand, some computational methods have applications in the field of neuroscience [3]. However, it is still a challenge to replicate the brain’s capabilities to solve complex visual problems (e.g. classification of stimuli) in AI. AI and neuroscience have attempted to address some common questions. One question that has a very important impact on both fields is the exact mechanisms of learning. There are many different observed learning capabilities, and in animals the mechanism of very simple learning paradigms like conditioning is still not fully known [4, 5]. One of the amazing capabilities of the human brain is generalization from past experiences, which is related to the degree of similarity between prior experience and novel situations [6]. Current artificial intelligence (AI) uses relatively simple and uniform network structures and rely on learning algorithms using large sets of training data. Neurons in artificial neural networks perform simplified integration of inputs and elicit spikes in response to super-threshold stimuli. In contrast, animal brains often accomplish complex learning tasks with limited training data. These observations have motivated researchers to believe that animals’ brain perform mainly unsupervised learning [7].

An artificial neural network is a rough analogy of synapses as matrices of numbers that change through training. However, synapses are complex machineries that perform complicated computations and interact with their neighbors in dynamic patterns [8]. Deep networks architectures as brain-inspired models of cortical circuits are successive layers of units named neurons and connected by synapses. These architectures have demonstrated capabilities in performing computer vision and speech recognition [9, 10]. The key mechanism of deep networks is the adjustment of the synapses to produce the desired outputs by providing training examples.

The important question is whether such simplified neural networks compared with biological circuits are sufficient to capture human-like learning and cognition in computers. Human learning may result from interactions between complex functional and structural elements in neurons and biological neural networks [10, 11]. Therefore, translating biological intelligence into algorithmic constructions may play a vital role in developing more efficient AI. For a survey on interaction between AI and neuroscience fields and advances in neuroscience-inspired AI like memory and learning systems we refer the readers to [12].

Classification algorithms, as one of the fundamental approaches in modern AI, assign data to categories by analyzing sets of training data. To evaluate performance of classification algorithms, some databases are being used. One of them is MNIST database that is a large database of handwritten digits that is commonly used for training various classification methods. MNIST database contains 60,000 training images and 10,000 testing images [13]. The classification methods that have used MNIST as a benchmark database are categorized into the following: Convolutional Neural Networks (CNN), Recurrent Neural Networks (RNN), Support Vector Machine (SVM) and Boltzmann machines. These classifiers vary in their recognition speed and accuracy of classification. Neural networks have demonstrated excellent performance for handwritten digit classification by extracting features from images in different ways. CNNs are the most important deep learning methods used in handwritten digit recognition with high accuracy, however, the execution time of the CNN methods are high compared to other neural networks based classification methods. All these methods requires optimization of large number of hyper-parameters of the network that should be optimized through training phase. For further reading on different methods that has used MNIST and their performance we refer to [14, 15].

Indeed, humans learn to classify objects by recognizing horizontal, vertical, orthogonal lines and circles composed handwritten digits. Some methods have considered these cognition-based feature selection approach [16, 17].

Although convolutional neural networks models attempt to extract features by learning hierarchical features, they are considered as ‘Black-Boxes’ [18, 19]. Here, Black-box convey the concept of returning the results of a decision task (e.g., classification) by artificial systems without providing sufficient details about its internal behavior or how to improve the internal mechanism underlying the decision. Therefore, the development of users-understandable AI systems can provide practical reasons to users and developers to explain the black-box models as much as possible. Such researches are named ‘Explainable AI (XAI)’ [20]. The technical explanation about an AI method may be appropriate for a data scientist as well as for researchers that need to apply an artificial system in analyzing their data.

Another important difference between biological neurons and their simplified models used in neural networks-based learning techniques is the assumed location of learned features throughout the learning process. Prior existing methods of artificial neural network based learning have been developed using the concept of synaptic weight modulation which is inspired by activity-dependent synaptic modulation. However, microtubules as cytoskeletal polymers inside biological neurons, are involved in learning and memory formation [21, 22]. One of such known mechanisms of memory formation by microtubules is its role in transport of MNDA receptors to synapses.

### Current study

In this work, we have developed an explainable feature extraction of handwritten digits method, which we applied to the MNIST dataset. The model is a single layer composed of 10 computational units (neurons) that encode the features of a handwritten digit and store them inside neurons instead of in synapses. In the next sections the methods is described and the basic results are presented. The importance of the method and some possibilities of developing and applying in different ways that are presented in the discussion.

## Methods

In this work, we assume that the human visual cortex detect horizontal, vertical and orthogonal lines and circles such that their combination leads to the retrieval of information to assign images into digit classes. The main challenge is an efficient way to convert such information into machine readable algorithm. In CNN approaches, such information is encoded in the synapses of connected neural layers via learning algorithms and is performed in special architecture of connected layers. In this work, features are extracted by single neurons and are stored inside the neurons as tensors.

For such proposed feature extraction method, a set of images with determined number of samples for each digit class is extracted randomly from MNIST database (includes 60,000 training images and 10,000 test images with digit class labels). The size of the training set of images per digit has in scale of single sample to higher values up to 1000 samples per digit.

Each image is presented as a 28*28 matrix of pixels. The preprocessing of data includes setting the value of pixels lower than 0.1 into zero. This preprocessing stage is required for efficient discrimination of lines that compose the images. Each training image is divided into eight region defined as I to VIII that are in different directions (**Figure. 1A**).

**Fig. 1.**
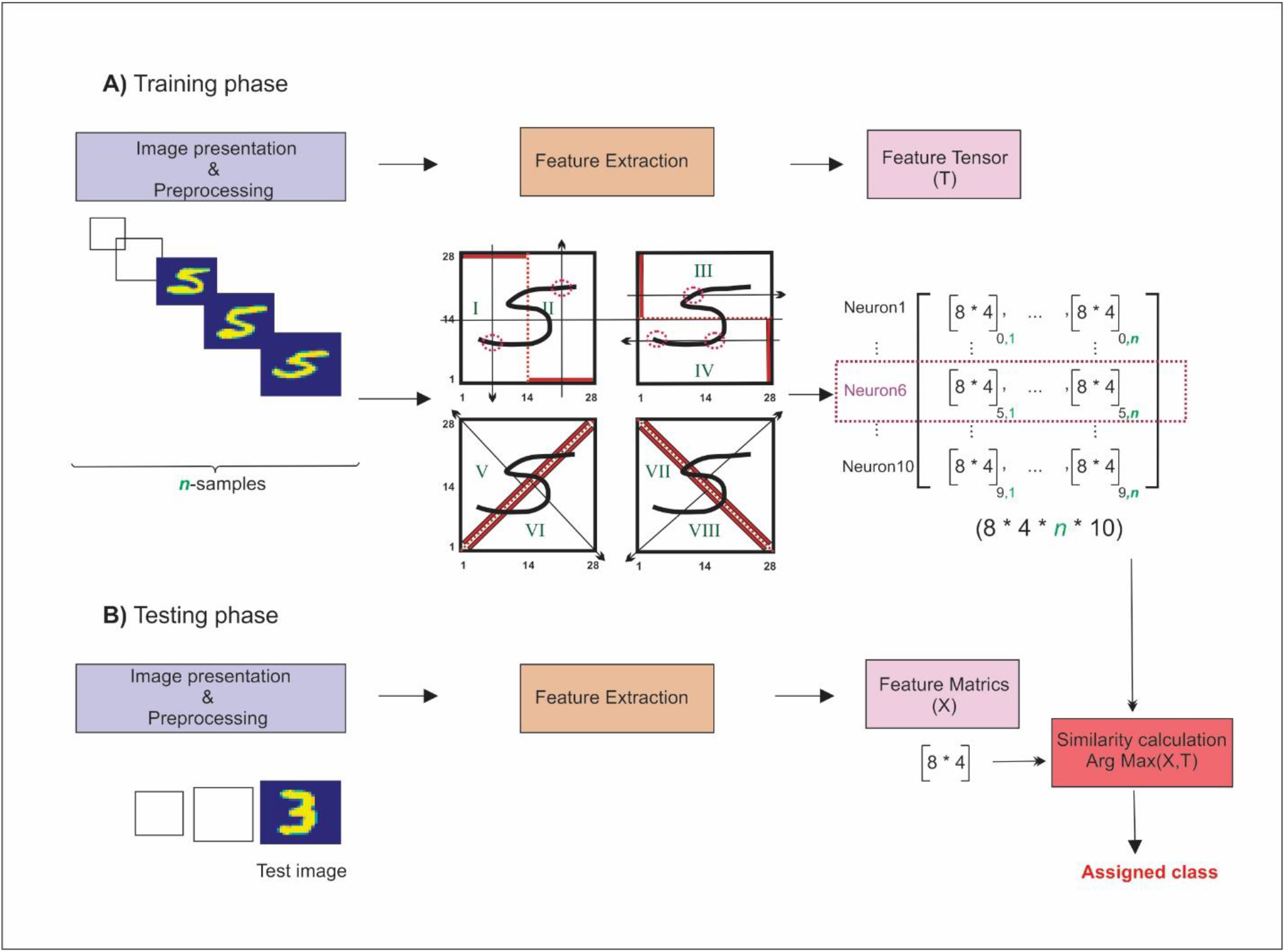
Model’s Structure. **A)** Training phase. Sets of training images with different size between one to 1000 samples per digit is selected from MNIST dataset. Each image is preprocessed by deleting the pixels with values lower than 0.1 (and setting them to 0.0). For feature extraction, the image that has a size of 28 *28 pixels are divided into 8 regions showing by I to VIII. In each region, one neuron scans the pixels and finds how many times their values change from zero to non-zero value and then to zero again and then information are stored in each neuron. The number of such changes between 1 and 4 is stored in 8*4 matrices. The total feature information on digits 0 to 9 in 10 neurons is stored as feature tensor for each digit class (**T**). **B)** Test phase. The features of the image is selected in a similar paradigm for each test image. The extracted feature matrix is constructed and then the similarity between the feature matrix (**X**) and T is calculated and the maximum value is used to assign the test image to one digit class.

To extract features from an image, one neuron scans a region pixel by pixel and detects the number of times pixels value is changed from zero to non-zero value and again to zero value. These numbers are counted and stored in a vector of size 4 (corresponding to one, two, three, and four times). Hence, for each image, the dimension of an image is reduced into a matrix of size 8*4. The information of training samples (*n* samples) are stored in a tensor of size 8*4\**n*. The process is done for all digits and eventually features tensor (or training information) is calculated and stored as a feature tensor (T) with size of 8*4\**n* *10, shown in **Figure. 1A**.

In the testing phase, each image of testing dataset that contains 10,000 images of random digits from zero to nine are preprocessed then its features are extracted using the proposed approach used for extracting training images features (**Figure. 1B**). After extracting the feature of the image (*X*), the cosine similarity between X and each element of T is calculated (**Equation. 1**). The maximum similarity is selected and the corresponding digit class is assigned to the given testing image.

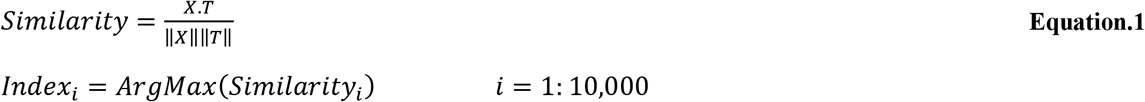

In order to evaluate the accuracy of classification of hand-written digits, different numbers of training samples per digit (*n*) is selected and images are chosen randomly from 60,000 training samples and the accuracy is measured. For each *n* value, 20 times the experiments are performed and the average accuracy is measured. We used *n* values as one random sample per digit to 50 samples per digit. In the next step, samples with higher values (from 100 to 1000) are selected to evaluate the enhancement of the efficiency as a function of the training size (**Figure 2**.).

**Fig. 2.**
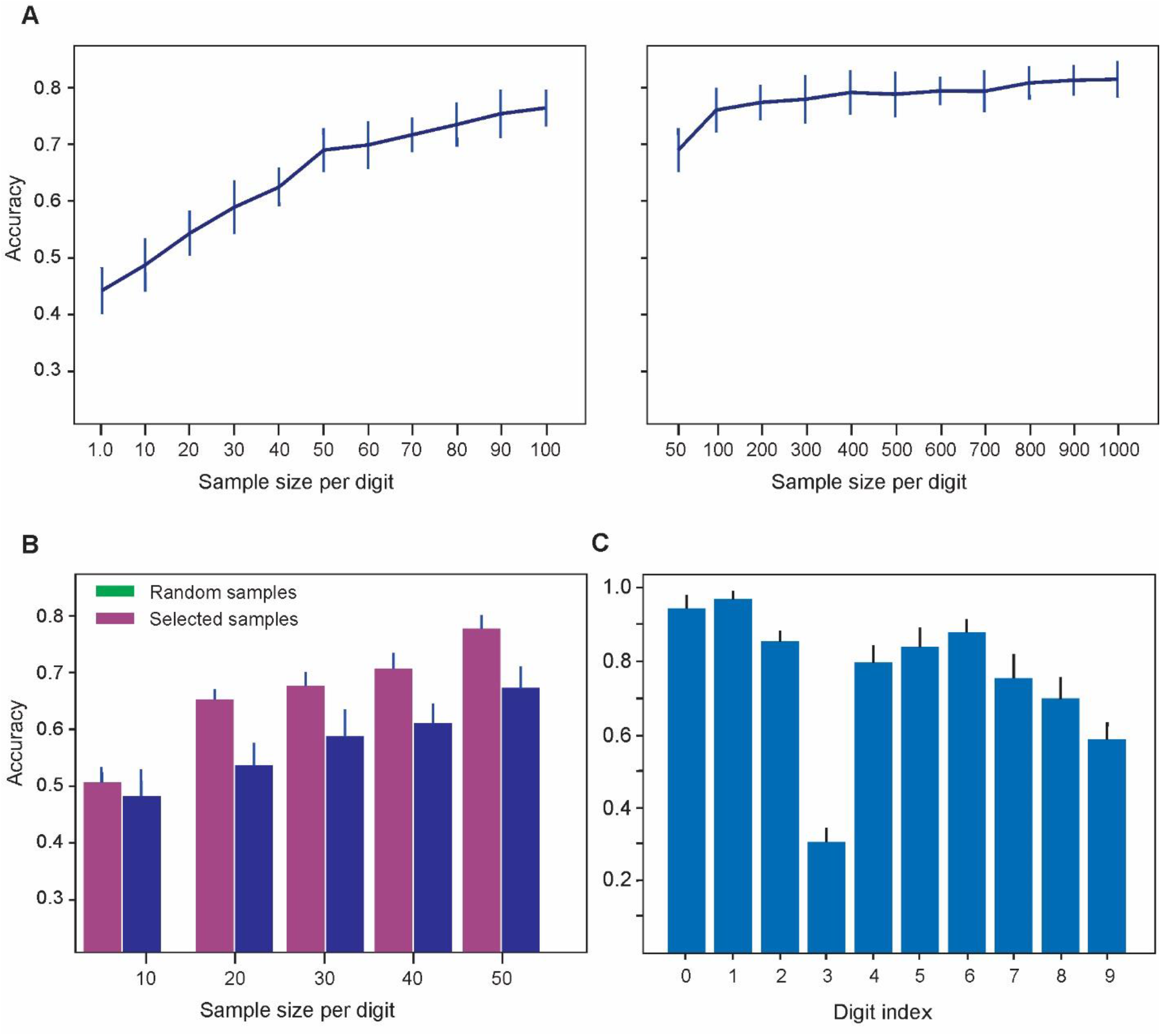
**A.** Accuracy of the method to assign digits class of images from the test set. An increase in the number of training sample per digit leads to enhancement of accuracy. The highest accuracy as 74 % in the range of 1 to 100 was obtained using 100 samples per digit. **B.** For each digit class *n* samples similar to canonical form of digits selected. **C.** Accuracy of the classification method to assign digit class of the test images for individual digits class (number of training sample per digit = 100). The method show lowest accuracy in assigning class of images from ‘3’. These results demonstrate the efficiency of the algorithm to extract features from the images except ‘3’.

In addition, to demonstrate the statistics of error for each digit class, correctly assigned digits of each class are also calculated for *n* equal to 100 (**Figure 3**.). One interesting question is the distribution of assigned images for each digit class to study the incorrect assigned class of each digit class. It helps interpreting the feature extraction algorithm to assign images to digits classes.

**Fig. 3.**
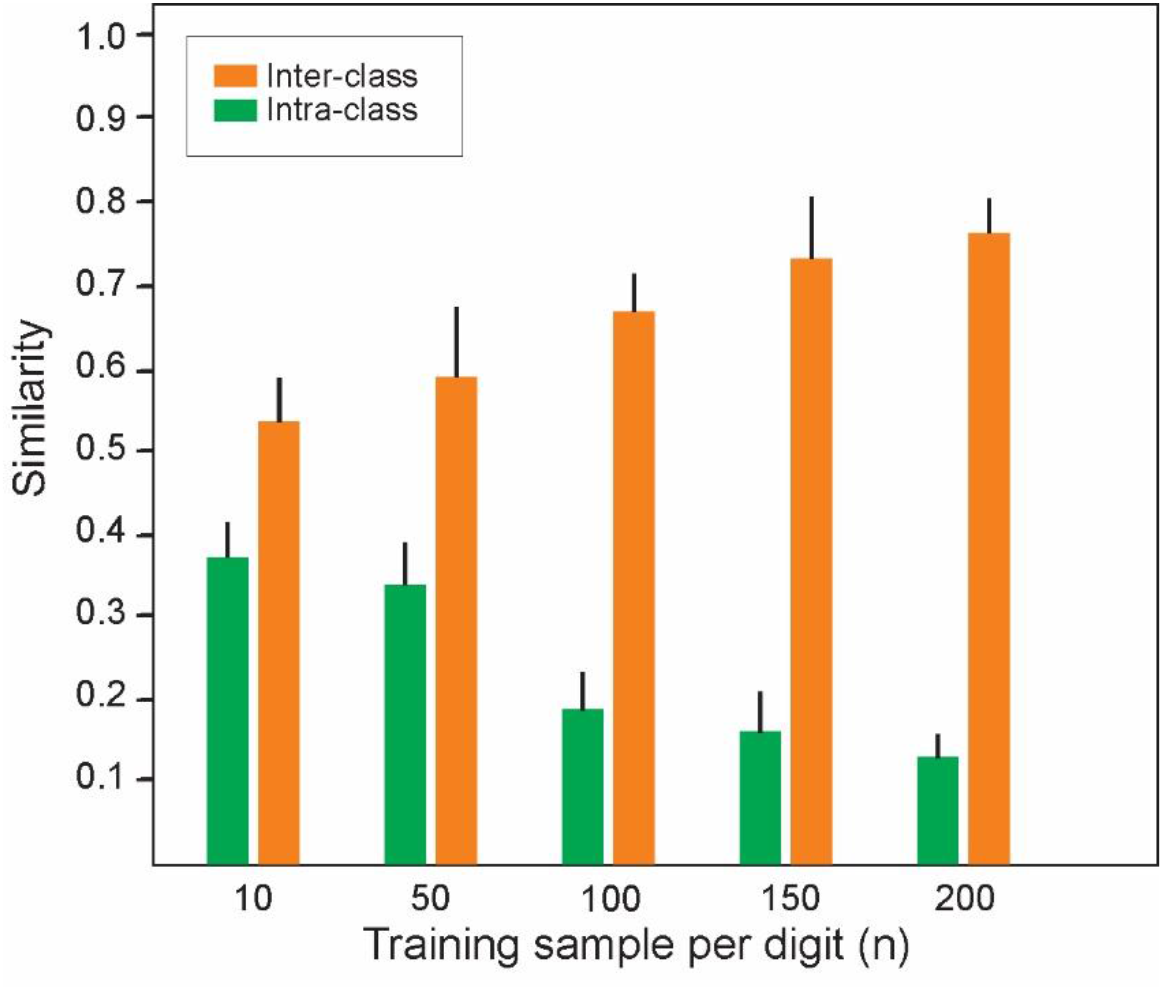
Average similarity of features matrices in training samples (n=100) of digit classes (inter-class) and between feature matrices of different digit classes (intra-class). Increase in number of training samples per digit leads to increase in inter-class similarity of feature matrices while it results in decrease in similarity of features matrices in intra-class.

## Experiments and Results

The feature extraction method trained on training images dataset of MNIST and then tested on test dataset of MNIST dataset. To evaluate accuracy of the method to assign images to proper digit classes, different number of training samples per digit (*n*) were applied as one parameter of the model.

**Figure 2.A** demonstrates the average accuracy of classification for different *n* values. The results show that an increase in *n*-value leads to enhancement of the classification accuracy. The increase of classification accuracy shows slow rate for *n*-values higher than 100 that is about 0.75 for n=100 and about 0.78 for n=1000. Using the whole training dataset, the system shows 86% classification accuracy. In these experiments for each digit class *n* samples are selected randomly from the training set. In addition, for each digit class *n* samples similar to canonical form of digits selected. **Figure 2.B** shows the enhancement of accuracy using ‘selected training samples’. **Figure 2.C** demonstrates the accuracy of the classification method to assign digit class of the test images for individual digits class (*n* = 100). The results show the lowest accuracy in assigning digit class of images for ‘3’. These results demonstrate the efficiency of the algorithm to extract features from the images except ‘3’.

The classification accuracy of method depends on high similarity of extracted features of n-training samples per digit while low similarity of clusters of samples for different digit classes. We calculated the similarity presented as cosine similarity measure of the feature tensors. **Figure .3** demonstrates average similarity for inter-class and intra-class samples for different *n*-values. The results show increase in average similarity in inter-class while decrease in intra-class average similarity of training samples for incremental *n* values.

Another important parameter of the proposed feature extraction method is the number of features that can be extracted from training samples of all digit classes. These features are shown in the Model Structure in **Figure .1**.

**Figure .4** shows accuracy of the classification method for number of samples equal to 100 for number of features used in the training and test phase of the experiments. Although, low number of regions leads to lower computational costs, the results demonstrate highest accuracy rate of the method for feature values equal to seven features and eight features. Therefore, in the presented results eight features were used.

**Fig. 4.**
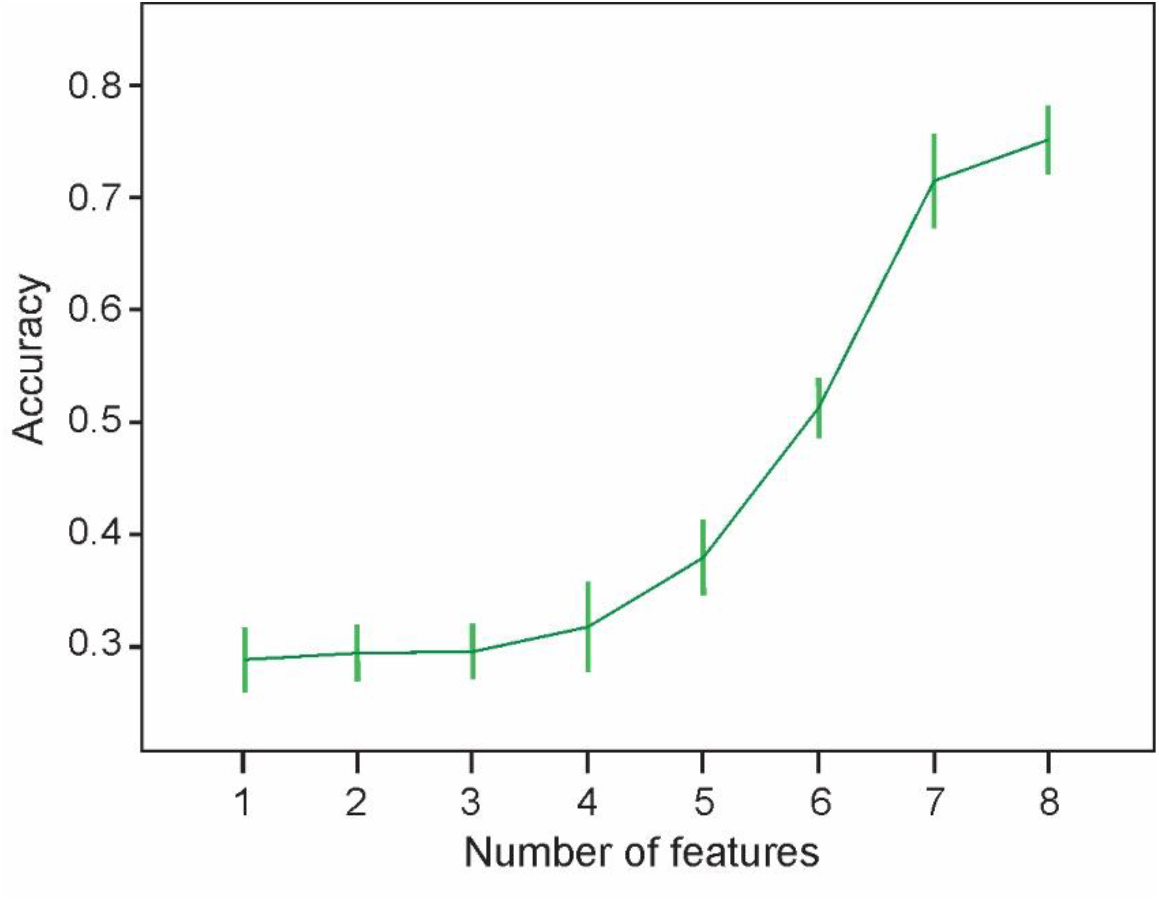
Accuracy of the classification method for different number of features (one to eight features) used in training and test phases. Increase in the feature numbers leads to enhancement in the accuracy. The number of training sample was set to 100. The highest accuracy is gained for seven and eight features.

In this work, we aim to study distribution of correct and incorrect assigned images of test dataset. These results are useful to evaluate the feature extraction algorithm to find misrecognized digit classes and to explain the possibilities to improve the method. **Figure .5A** shows the distribution of assigned classes (0 to 9) for test samples for *n* equal to 100. For digit classes with low accuracy it is important to find misrecognized digit classed. The results show that digit class ‘9’ is mainly misrecognized with digit class ‘4’. For the lowest accuracy belongs to digit class ‘3’ and shown by green box, the test samples are misrecognized with digit class ‘7’. Therefore, to improve the feature extraction method one need to concentrate the possibilities of proper encoding of such digits. Such distribution information provides the explanation to users and developers to find how to improve the classification model.

**Fig. 5.**
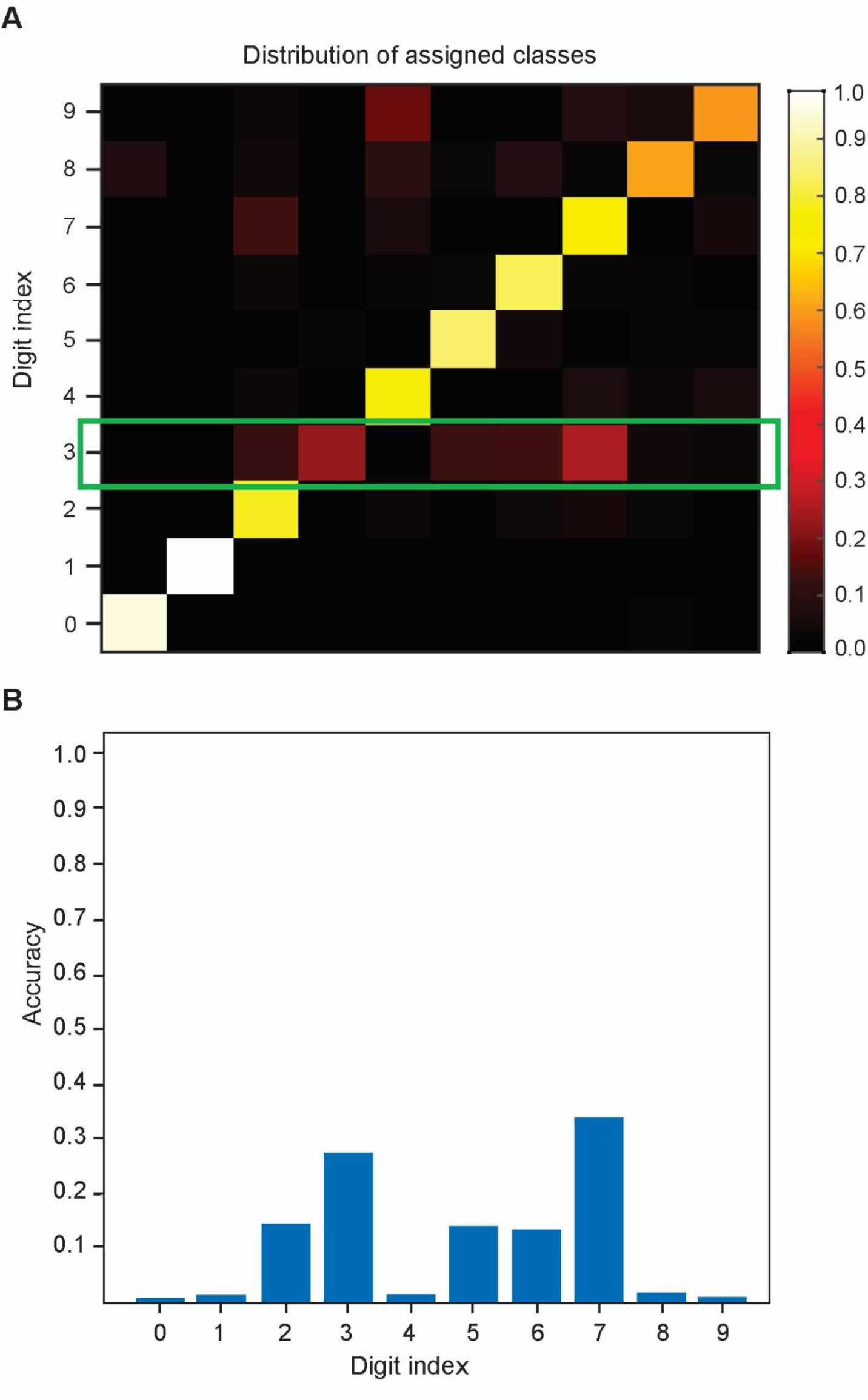
**A.** Distribution of assigned classes per digits. For the digit classes that the model has shown low accuracy the results show the distribution of assigning images to different classes. For example for digit class ‘3’ mainly the model has assigned incorrectly to digit class ‘7’. For digit class ‘9’ and ‘7’ incorrectly assigned class has been ‘4’ and ‘2’, respectively. **B.** Distribution of assigned test samples for digit ‘3’ shown in green box in **A.** The results shows that digit ‘3’ is mainly misrecognized with ‘7’.

## Discussion

Artificial intelligence research aims to mimic cognitive skills of animals and humans. Current AI methods are mostly based on statistical methods not on dynamical cellular and synaptic mechanisms [23]. One important class of modern AI method is deep neural networks methods that are mainly based on inspiration from the general function and architecture of biological neural systems.

In recent years, AI has focused on neuroscience-inspired artificial architectures that helps image and audio analysis problems [3]. In all these application for future AI, the main challenge is how to transform known biological mechanisms at synaptic and neural network levels into machine readable algorithms. In neural networks, algorithms are based on matrices and operation on matrices. However, in biological neural systems computations are performed in a continuous manner conducted by complex networks of molecules and synaptic channels.

Deep learning methods as images classification tools are often perceived as “black-boxes” as they are given inputs and produce desired outputs and, therefore, are unexplainable. There are many deep learning methods used MNIST dataset as a benchmark to evaluate their accuracy. In this work, we developed an interpretable neural network model that can explain the results. To our knowledge, it is the first attempt to extract handwritten digits features by non-synaptic memory in a single neural layer. Although the method use a single layer of neurons, it is possible to developed layers of such single computational units for classification and image processing problems as future works.

In human, digits are recognized by detecting horizontal, vertical, orthogonal lines and the circles that their combinations which construct digits. The proposed method present a novel class of computational unit to learn and detect these features. We named these units neurons as they can be connected to other units by synapses. In addition, these neurons instead of storing information in their synaptic weights store information inside the neurons that is inspired by biological evidences of existing such intrinsic mechanisms to memorize inputs pattern. The proposed manner of extracting and storing the information of images inside the neurons demonstrates the possibility of using single neurons as feature extracting units. In this work, for the first time accuracy of the classification of handwritten digits is calculated for individual digit classes. Hence, it is an explainable intelligent system for handwritten digit recognition. This method has revealed the lowest accuracy of classification for digit class ‘3’ as it is misrecognized mainly with digit ‘7’. It seems that the method is not efficient in encoding semi-circles that construct ‘3’. Therefore, one possible way of improving our work would be to enhance feature extraction algorithm for digit ‘3’ and ‘7’. In addition, our results demonstrate that our network model misrecognizes ‘9’ and ‘4’. Similarly, it seems that the method is not efficient in encoding circle that construct digit ‘9’ and so misrecognize it with lines constructing digit ‘4’. A cognitive experimental study on reading handwritten digits has shown that ‘9’ and ‘4’ are misinterpreted by many people because of similarity in their appearance [24]. Thus, our results are in agreement with these experimental findings.

The results show that an increase in number of training samples per digit leads to enhancement of inter-class similarity of features matrices to enrich encoding of training data while decrease the similarity of intra-class similarity of features matrices. Therefore, an increase in size of the training sample per digit plays an important role in enhancement of the method.

The idea of feature of images as vertical and horizontal direction of written digits for hand-written digits classification has been already presented [16]. In this work, mathematical morphology to exploit the shape of the ten digits in in MNIST database is used. Morphology was used to separate digits into two groups: (a) Groups with blobs with/without stems {0, 4, 6, 8, 9}; and (b) Groups with stems {1, 2, 3, 5, 7}. A concept called ‘connected component’ was defined in this work to identify the digits with blobs and stems.

Another method developed a Support Vector Machine based handwritten digit recognition system for Arabic numerals that are very different from Latin digits in shape. The method is based on transition information in the vertical and horizontal directions of the images [17]. They have used a different features extraction method called chain code histogram (CCH) with SVM as the classifier. This method reduces the dimensionality of the feature space without decreasing the performance of the classifier. The results showed that the digits with similar shapes are difficult to be recognized by the system and even for human being.

In this work, we extract such features in a very different way, one neurons scan eight defined regions of the image in different direction and count the number of times they cross a line (from 1 to 4). Such feature extraction method presents a new explainable image feature extraction method that can be developed in deep neural networks techniques.

## Conclusions and future work

Here, we proposed a new feature extraction method for handwritten digit classification that is based on storing image information inside the 10 single neurons as non-synaptic memory. The method extracts image features as lines, circle and semi-circles. The method allows to compare the misrecognized images for each digit class and hence allows explaining how the images have misrecognized. It provide possibilities to improve the classification algorithm according to the results. The proposed method is based on single neurons as encoding units. However, it can be developed as networks of such units to explore the application of the method for other image classification problems.

## References

[1] Wang, Yingxu, Sam Kwong, Henry Leung, Jianhua Lu, Michael H. Smith, Ljiljana Trajkovic, Edward Tunstel, Konstantinos N. Plataniotis, Gary G. Yen, and Witold Kinsner. “Brain-inspired systems: A transdisciplinary exploration on cognitive cybernetics, humanity, and systems science toward autonomous artificial intelligence.” IEEE Systems, Man, and Cybernetics Magazine 6, no. 1 (2020): 6–13.

[2] Ullman, Shimon. “Using neuroscience to develop artificial intelligence.” Science 363, no. 6428 (2019): 692–693.

[3] Hassabis, Demis, Dharshan Kumaran, Christopher Summerfield, and Matthew Botvinick. “Neuroscience-inspired artificial intelligence.” Neuron 95, no. 2 (2017): 245–258.

[4] Faghihi, Faramarz, Ahmed A. Moustafa, Ralf Heinrich, and Florentin Wörgötter. “A computational model of conditioning inspired by Drosophila olfactory system.” Neural Networks 87 (2017): 96–108.

[5] Delahunt, Charles B., Jeffrey A. Riffell, and J. Nathan Kutz. “Biological mechanisms for learning: a computational model of olfactory learning in the Manduca sexta moth, with applications to neural nets.” Frontiers in computational neuroscience 12 (2018): 102.

[6] FeldmanHall, Oriel, Joseph E. Dunsmoor, Alexa Tompary, Lindsay E. Hunter, Alexander Todorov, and Elizabeth A. Phelps. “Stimulus generalization as a mechanism for learning to trust.” Proceedings of the National Academy of Sciences 115, no. 7 (2018): E1690–E1697.

[7] Zador, Anthony M. “A critique of pure learning and what artificial neural networks can learn from animal brains.” Nature communications 10, no. 1 (2019): 1–7.

[8] LeCun, Yann, Yoshua Bengio, and Geoffrey Hinton. “Deep learning.” nature 521, no. 7553 (2015): 436–444.

[9] Voulodimos, Athanasios, Nikolaos Doulamis, Anastasios Doulamis, and Eftychios Protopapadakis. “Deep learning for computer vision: A brief review.” Computational intelligence and neuroscience 2018 (2018).

[10] Deng, Li, and John C. Platt. “Ensemble deep learning for speech recognition.” In Fifteenth Annual Conference of the International Speech Communication Association. 2014.

[11] Faghihi, Faramarz, and Ahmed A. Moustafa. “Combined computational systems biology and computational neuroscience approaches help develop of future “cognitive developmental robotics”.” Frontiers in neurorobotics 11 (2017): 63.

[12] Li, Junjun, Zhijun Li, Fei Chen, Antonio Bicchi, Yu Sun, and Toshio Fukuda. “Combined sensing, cognition, learning, and control for developing future neuro-robotics systems: a survey.” IEEE Transactions on Cognitive and Developmental Systems 11, no. 2 (2019): 148–161.

[13] Kussul, Ernst, and Tatiana Baidyk. “Improved method of handwritten digit recognition tested on MNIST database.” Image and Vision Computing 22, no. 12 (2004): 971–981.

[14] Albahli, Saleh, Fatimah Alhassan, Waleed Albattah, and Rehan Ullah Khan. “Handwritten Digit Recognition: Hyperparameters-Based Analysis.” Applied Sciences 10, no. 17 (2020): 5988.

[15] Sethi, Rohan, and Ila Kaushik. “Hand Written Digit Recognition using Machine Learning.” In 2020 IEEE 9th International Conference on Communication Systems and Network Technologies (CSNT), pp. 49–54. IEEE, 2020.

[16] Kumar, V. Vijaya, A. Srikrishna, B. Raveendra Babu, and M. Radhika Mani. “Classification and recognition of handwritten digits by using mathematical morphology.” Sadhana 35, no. 4 (2010): 419–426.

[17] Boukharouba, Abdelhak, and Abdelhak Bennia. “Novel feature extraction technique for the recognition of handwritten digits.” Applied Computing and Informatics 13, no. 1 (2017): 19–26.

[18] Zhao, Hui-huang, and Han Liu. “Multiple classifiers fusion and CNN feature extraction for handwritten digits recognition.” Granular Computing 5, no. 3 (2020): 411–418.

[19] Chaki, Jyotismita, and Nilanjan Dey. “Fragmented handwritten digit recognition using grading scheme and fuzzy rules.” Sādhanā 45, no. 1 (2020): 1–23.

[20] Guidotti, Riccardo, Anna Monreale, Salvatore Ruggieri, Franco Turini, Fosca Giannotti, and Dino Pedreschi. “A survey of methods for explaining black box models.” ACM computing surveys (CSUR) 51, no. 5 (2018): 1–42.

[21] Martel, Guillaume, Shusaku Uchida, Charles Hevi, Itzamarie Chévere-Torres, Ileana Fuentes, Young Jin Park, Hannah Hafeez, Hirotaka Yamagata, Yoshifumi Watanabe, and Gleb P. Shumyatsky. “Genetic demonstration of a role for stathmin in adult hippocampal neurogenesis, spinogenesis, and NMDA receptor-dependent memory.” Journal of Neuroscience 36, no. 4 (2016): 1185–1202.

[22] Dent, Erik W. “Of microtubules and memory: implications for microtubule dynamics in dendrites and spines.” Molecular biology of the cell 28, no. 1 (2017): 1–8.

[23] Hastie, Trevor, Robert Tibshirani, and Jerome Friedman. “An introduction to statistical learning.” (2009).

[24] Grissinger, Matthew. “Misidentification of Alphanumeric Symbols Plays a Role in Errors.” Pharmacy and Therapeutics 42, no. 10 (2017): 604.

